# AistSeq: An In-House Tn5-Based Plasmid Sequencing Platform Using A Compact Benchtop Sequencer

**DOI:** 10.1101/2024.11.04.618112

**Authors:** Hayato Suzuki, Jutapat Romsuk, Akiyoshi Nakamura, Kosuke Moriwaki, Shigeo S. Sugano

**Affiliations:** Plant Molecular Technology Research Group, Bioproduction Research Institute, The National Institute of Advanced Industrial Science and Technology (AIST), Sapporo, Japan; Plant Gene Regulation Research Group, Bioproduction Research Institute, The National Institute of Advanced Industrial Science and Technology (AIST), Tsukuba, Japan

**Keywords:** Tn5 transposase purification, next-generation sequencing (NGS), plasmid sequencing, pipeline, iSeq100

## Abstract

Sequence verification of plasmids is a fundamental process in synthetic biology. For plasmid sequence verification using next-generation sequencing (NGS) library preparation, Tn5 transposase is widely used. Streamlined sequencing workflow for laboratory-scale applications is important; however, recombinant Tn5 production *in-house* can be laborious. In this study, we demonstrated that the addition of a large soluble tag was not essential for purification and that the fusion of a His10 tag and protein A was sufficient to purify sufficient amounts of active Tn5 transposase. In addition, we evaluated exonuclease-based genomic DNA digestion for plasmid sequencing from an *E. coli* lysate and the data analysis pipeline of sequences derived from the Illumina iSeq100 platform for *de novo* assembly, reference mapping, and annotation. This study proposes a simple workflow of <u>a</u>n in-hou<u>s</u>e <u>T</u>n5-based plasmid <u>Seq</u>uencing platform using a compact benchtop sequencer (AistSeq).

**Graphical abstract:** 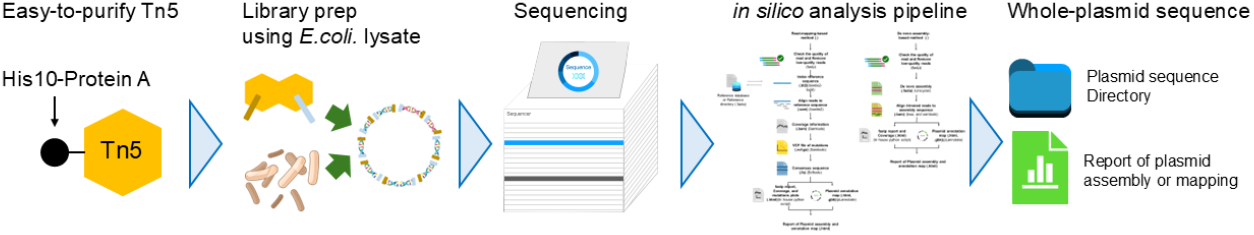

## Introduction

Tn5 transposase, discovered in the early 1990s in *Escherichia coli*^1^, is pivotal for next-generation sequencing (NGS) library preparation. It forms a ternary complex known as the Tn5 transposome, which consists of a homodimer of the Tn5 protein and its DNA cargo. This offers versatility in custom DNA and adapter payloads^2^. The Tn5 transposome can fragment DNAs and attach sequencing adapters simultaneously, thus streamlining NGS workflows, such as the Nextera library preparation kit^3^ and ATAC-Seq^4^. Recent studies have demonstrated the ability of Tn5 to target DNA/RNA hybrids, thereby broadening its applications to RNA-seq^5^; however, recombinant Tn5 transposase is not so easy to produce in the laboratory scale. There are several methods to obtaining enough purified Tn5, including TnY^6,7,8^, however, the fusion of large soluble tags or chitin-binding protein tags to the transposons is necessary. Recently, OpenTn5 open-source and easy-to-purify Tn5 was proposed^9^, applicability of this methodology has not been explored.

These developments have addressed some of the challenges faced by industrial-scale synthetic biologists, such as the high costs associated with Sanger sequencing for plasmid verification. Integrating Tn5 into plasmid sequencing workflows drastically reduces costs, thus enabling sequencing of over 4,000 plasmids in a single Illumina MiSeq run for <$3 sequencing each^10,11^. To minimize preparation costs, plasmid sequencing from an *E. coli* lysate without purifications has been demonstrated^11^. In addition, various software programs and pipelines have been developed to support and verify plasmid sequencing analysis^11^. Likewise, there is a significant demand for streamlined plasmid sequencing methods utilizing Tn5 transposome. Here, we propose a new approach to Tn5 transposase purification, in which only a His10 tag and protein A (pA) were used to purify Tn5 transposase. In addition, we developed a simple in-house workflow for plasmid-DNA sequencing of *E. coli*. lysates by Illumina iSeq100 platform. We also established an alternative bioinformatics pipeline to accommodate *de novo* sequence assembly and alignment with reference plasmid sequences for generating a FASTA file comprising the sequence of the plasmid and an annotated map, encompassing insertion, point mutations, and deletions to the reference. This streamlined workflow for the verification of plasmid-DNA sequences would contribute to molecular biology including the synthetic biology field.

## Results and Discussion

### Expression of His10-pA-Tn5 is sufficient to purify abundant Tn5 proteins

Recombinant Tn5 transposase tightly binds to nucleic acids during the purification step^3,4,7^. Although a chitin column was used for the purification of Tn5 transposase, a recent report revealed that GB1-Tn5 transposase with a His10-tag can be purified without any nucleic acids by washing with a high salt buffer (800 mM NaCl) on a Ni-affinity column^7^. We constructed a His10-tagged pA-Tn5 expression vector, which is a smaller tag compared with GB1, and purified recombinant pA-Tn5 using a Ni-affinity column. Recently reported OpenTn5 method is like our approach that it takes advantage of protein G instead of pA^9^.

Following Ni-affinity purification, recombinant pA-Tn5 transposase was further purified by size exclusion chromatography. The peaks from size exclusion chromatography (Figure 1a) and SDS-PAGE analysis of the eluted fractions (Figure 1b) suggested that the dimer fractions of pA-Tn5 transposase did not contain any nucleic acids, which was confirmed by measuring the 260 nm/280 nm ratio. The dimer fractions containing pA-Tn5 transposase were collected and concentrated to 60 μM. The purified pA-Tn5 assembled to oligonucleotides (harboring mosaic end sequences with Illumina adapter sequences) successfully fragmented pUC18 plasmids, like commercially available Tn5 (Figure 1c), which was also supported by the results of iSeq100 sequencing (Figure 1d).

**Figure 1.**
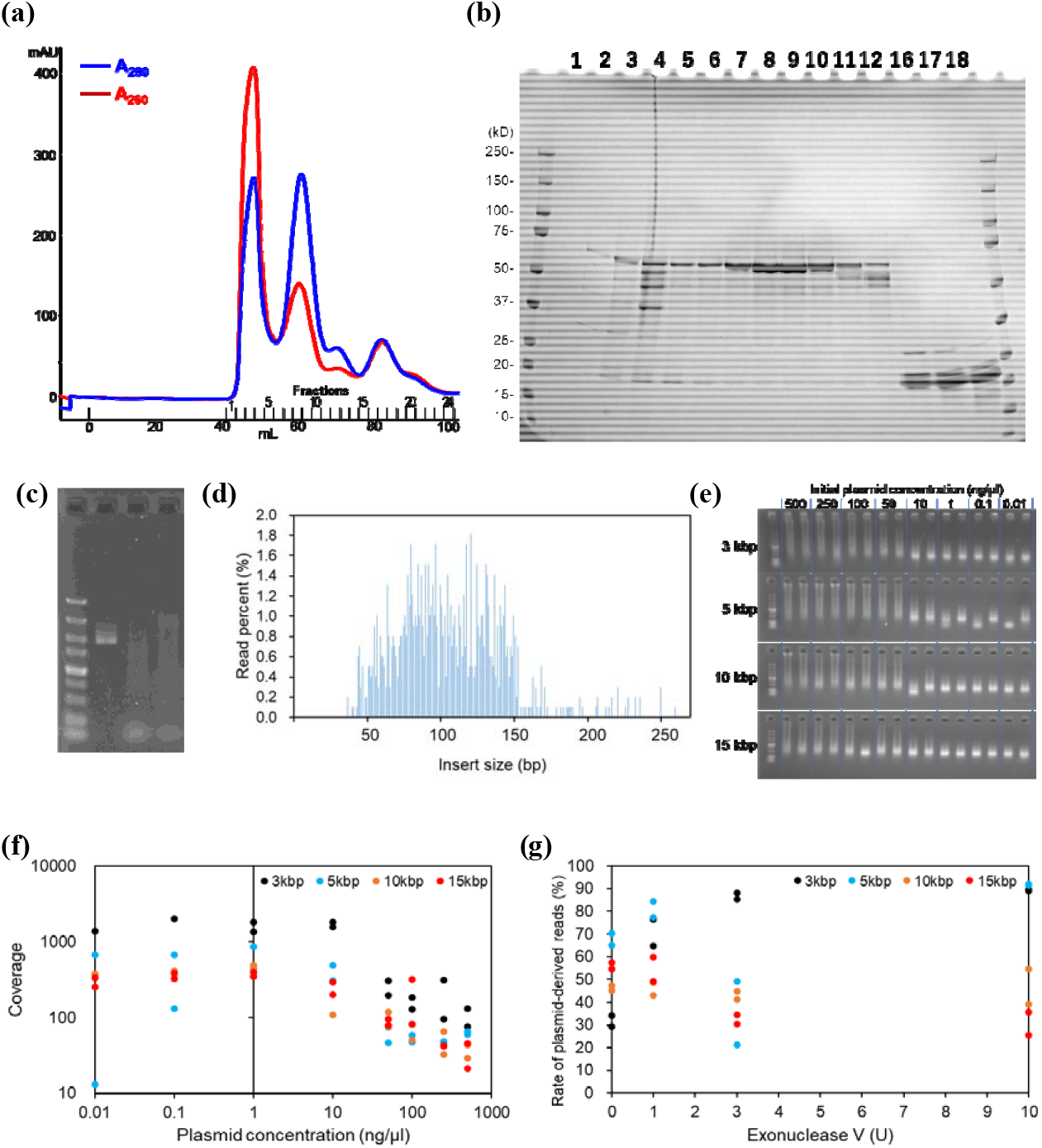
Tn5 transposon purification, Tn5-based plasmid digestion, and output of NGS analysis of plasmid prepared by several methodologies. Size exclusion chromatogram **(a)** and SDS-PAGE to evaluate Tn5 transposase purification. **(c)** Gel electrophoresis of intact pUC18 plasmid (left), digested by commercial Tn5 (diagenode; middle) and digested by in-house His10-pA-Tn5 (right). **(d)** Size distribution of sequencing inserts confirmed by iSeq100 sequencing. **(e)** Tn5 digestion of plasmids with different lengths and concentrations. **(f)** Sequencing coverage plots for the NGS analysis of plasmids with different lengths and concentrations. **(g)** The ratio of plasmid-derived reads to total reads in the libraries constructed from the supernatant of *E. coli* lysates incubated with various concentrations of ExoV.

### Tn5-based whole plasmid sequencing and the plasmid size/concentration dependency

Tagmentation of 3, 5, 10, and 15 kbp plasmids at concentrations ranging from 0.01 to 500 ng/μL was evaluated using in-house-prepared pA-Tn5. Gel electrophoresis of the PCR products revealed that higher initial plasmid concentrations resulted in larger average DNA fragment sizes (Figure 1e). The sequencing coverage was lower at higher initial plasmid concentrations (Figure 1f), which was attributed to insufficient Tn5 proteins resulting from excess plasmid and predominantly larger DNA fragments. Thus, maintaining an appropriate Tn5: plasmid ratio is necessary for DNA library preparation. For larger plasmids (10 and 15 kbp), diluted plasmids (<10 ng/μL) are preferable.

### Sequencing plasmid from E. coli lysates and assessing genome contamination

Direct NGS library preparation of the plasmid to be sequenced from *E. coli* lysates offers time and cost savings; however, *E. coli* genome contamination can hinder plasmid-derived read recovery^11^. We evaluated exonuclease V (ExoV) efficacy in selectively digesting damaged genomes before library preparation because circular plasmids without ends are not digested by ExoV. ExoV (0, 1, 3, and 10 U)-treated lysates of *E. coli* harboring plasmids of various lengths (3, 5, 10, and 15 kb) were assessed after the separation of the lysate pellets and supernatants (Table S1). The lysates had higher genomic DNA content, which affected plasmid fragmentation and low read quality compared with the supernatants. Using the pellet samples, full-length plasmid sequences were obtained for 3 and 5 kb plasmids, but larger plasmids (10 and 15 kb) were incompletely assembled. Higher ExoV concentrations negatively affected sequencing results for larger plasmids (supernatant samples), which was likely caused by susceptibility to linearization of plasmid during vortexing and pipetting (Figure 1g). Remarkably, assembly into one circular sequence was successfully achieved in supernatant samples treated with 0 or 1 U of ExoV, whereas the pellet samples yielded few results through *de novo* assembly (Table S1). These findings suggest the universal suitability of supernatant samples treated with 0 or 1 U ExoV.

**Table S1.**
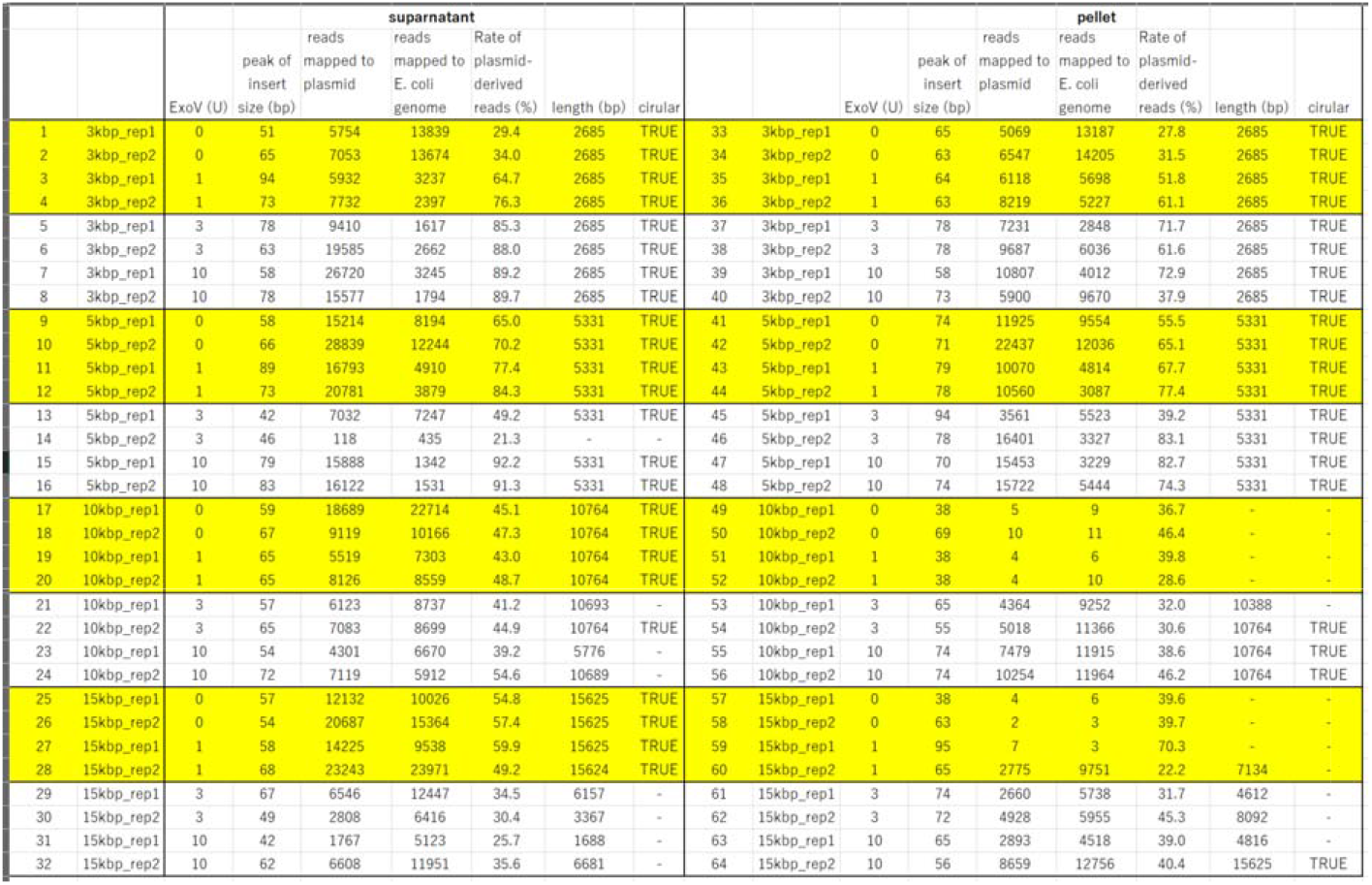
Summary of the results of plasmid sequencing from *E. coli* lysates.

### In silico Analysis Pipeline

As well as the wet experimental workflow, we developed a comprehensive bioinformatics pipeline for the analysis and reporting of plasmid sequencing obtained from iSeq100 paired-end 150 bp data (Figure 2). This program offers automating *de novo* assembly, mapping to a reference sequence, and showing metrics that described above (Table S1). *De novo* assembly of plasmid was conducted using Unicycler^11^, originally proposed as a *de novo* assembly pipeline for bacterial genomes. Mapping to the references was done by Bowtie2^13^. As described in Table S1, the developed pipeline yielded one circular sequence (successful assembly) for 44 out of 64 samples, whereas mapping to a reference was successfully achieved for 57 samples, except those with read numbers that were too small (<10 reads) (Table S1; Figure S1a, b). For mapping results, consensus sequences were outputted as “.fasta” files and were accompanied by coverage plots for each plasmid (Figure S1c). Mutated nucleotide positions were identified by bcftools^14^ and BEDtools^15^, including an A to T mutation at 1062 bp in the *ori* of the pUC18 plasmid, which was preserved in the samples tested (Figure S1d). The output consensus sequences were automatically annotated by pLannotate^16^ to enhance user-friendliness (Figure 2, Figure S1e). Combined with the simple pA-Tn5 purification, library preparation using *E. coli* lysate supernatant, and our analysis pipeline, we provide a streamlined plasmid verification workflow, <u>a</u>n in-hou<u>s</u>e <u>T</u>n5-based plasmid sequencing platform using a compact benchtop <u>seq</u>uencer, designated AistSeq.

**Figure 2.**
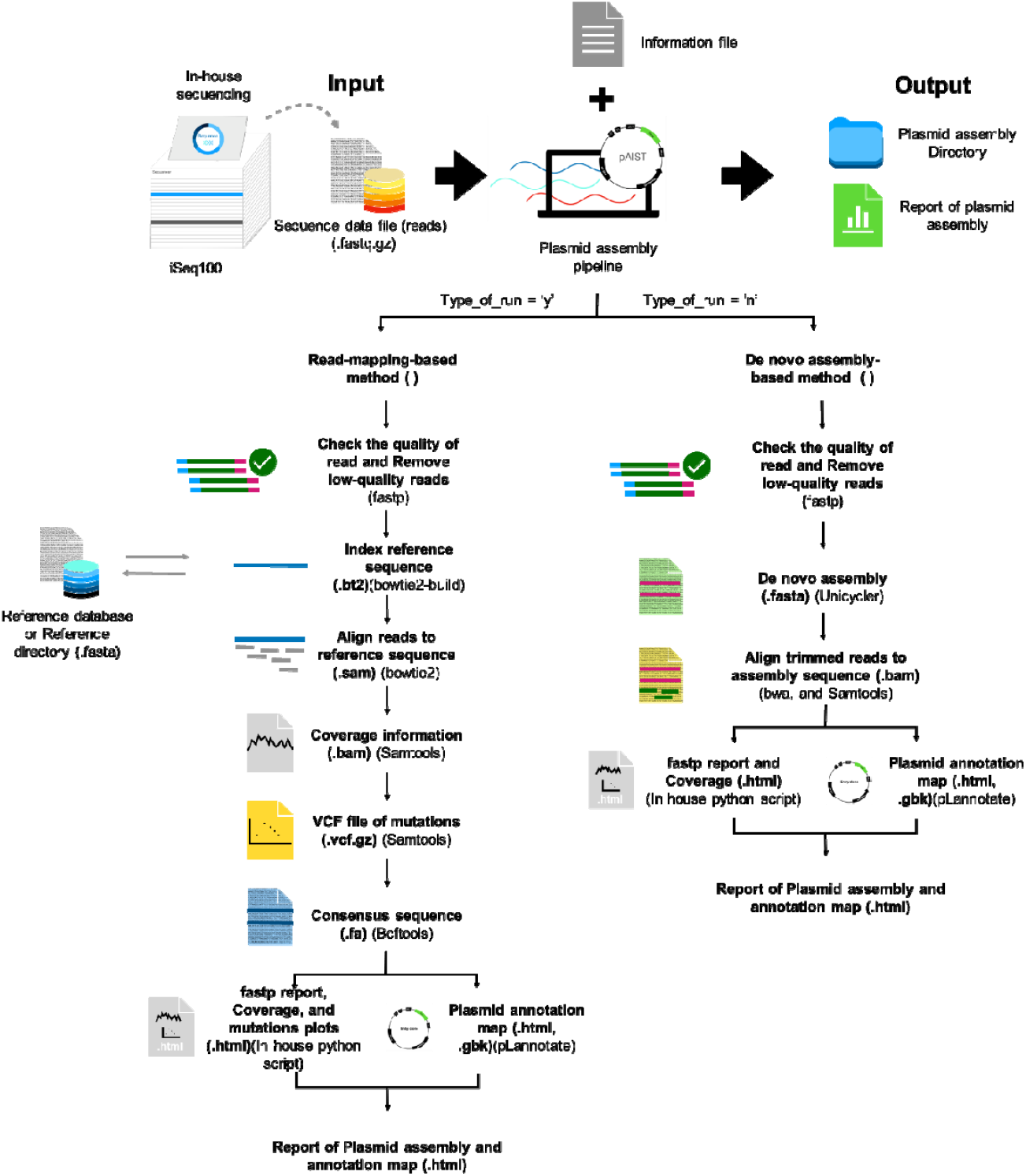
Workflow of the bioinformatic pipeline for plasmid sequencing developed in this study.

**Figure S1.**
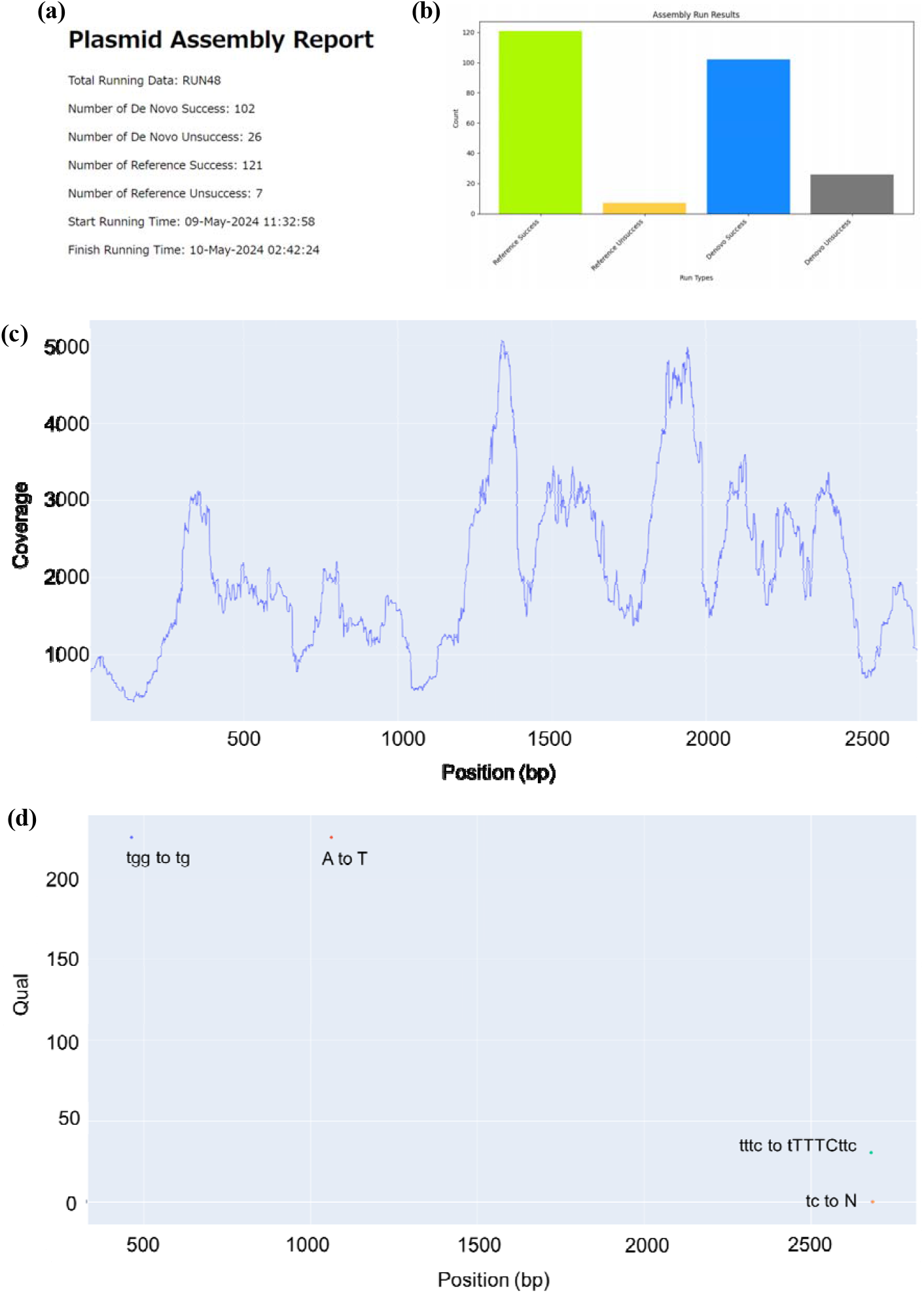
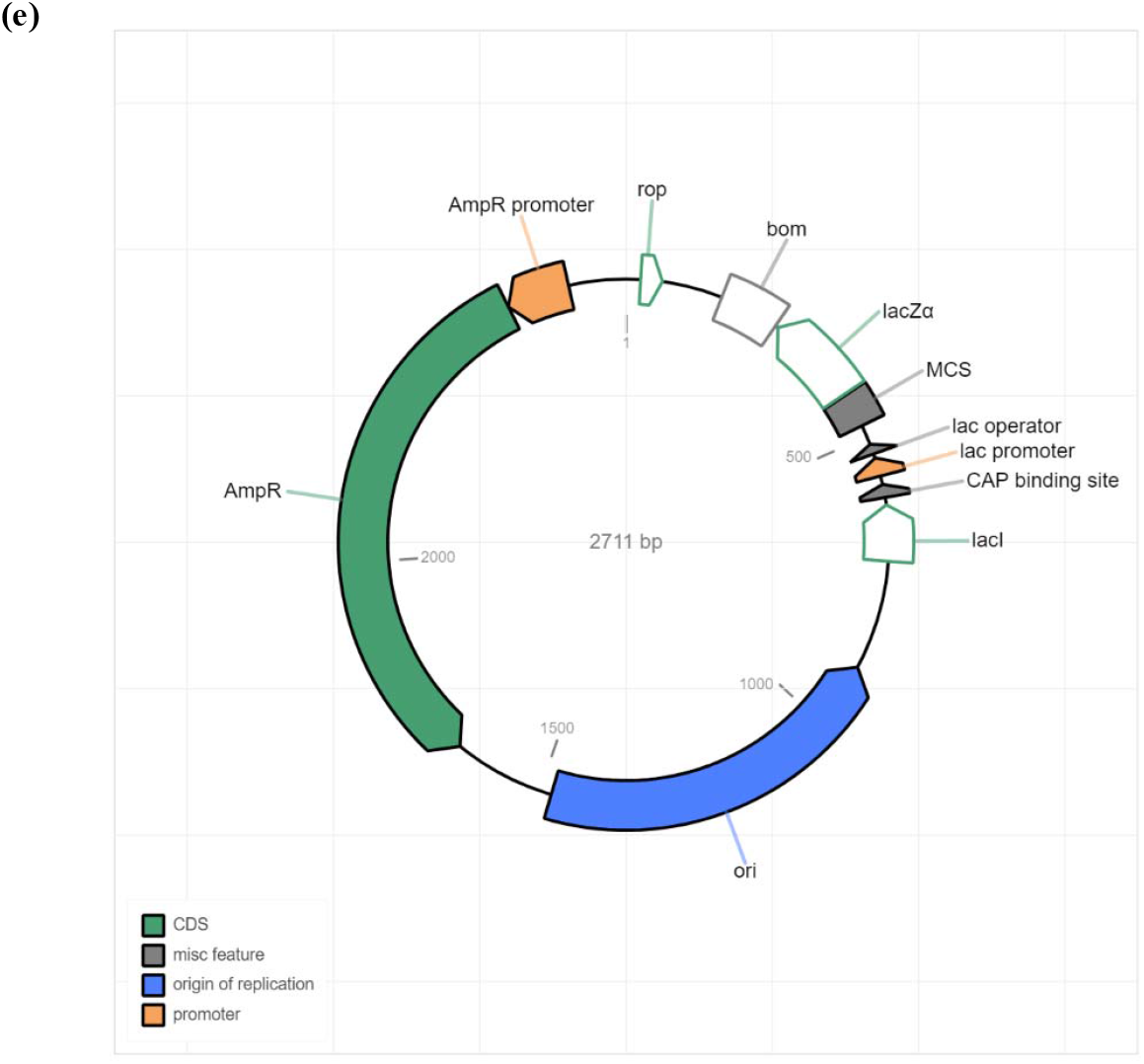
Output data of the pipeline. (a) Representative image of the report of mapping to a reference and *de novo* assembly. (b) Representative coverage plot of a sample with low sequencing coverage based on the data from Table S1. (c) Representative coverage plot of a sample with sufficient sequencing coverage based on the data from Table S1. (d) Representative mutation plot for the pUC18 plasmid analyzed in this study. (e) The Representative output of an annotated plasmid map.

## Materials and Methods

### Recombinant Tn5 preparation

The pA-Tn5 gene (Addgene plasmid #124601) was cloned into pET26b (Novagen) along with an N-terminal His10-tag. The plasmid was transformed into the *E. coli* strain BL21-AI (Invitrogen) for recombinant pA-Tn5 expression. Bacteria containing the plasmid were pre-cultured in a medium containing 25 μg/mL kanamycin. When the OD_600_ reached 0.6, 0.2% (w/v) arabinose was added to induce protein expression at 18°C overnight. Cell pellets were resuspended in buffer A [50 mM Tris-HCl, pH 7.5, 800 mM NaCl, 1 mM MgCl_2_, 1 mM DTT, 10% (v/v) glycerol, and 20 mM imidazole] with 0.5 mg/mL lysozyme and 0.1 mg/mL DNase I, then disrupted by sonication. The supernatant was loaded onto a 1 ml Ni-NTA Superflow column pre-equilibrated with buffer A. After loading the sample, the column was washed with buffer B [50 mM Tris-HCl, pH 7.5, 800 mM NaCl, 1 mM MgCl_2_, 1 mM DTT, 10% (v/v) glycerol, and 125 mM imidazole], and the recombinant His10-pA-Tn5 was eluted with 250 mM imidazole. The eluted sample was loaded onto a HiLoad 16/600 Superdex 200 pg column (Cytiva) in buffer C [50 mM Tris-HCl, pH 7.5, 800 mM NaCl, 0.2 mM EDTA, 2 mM DTT, 10% (v/v) glycerol]. The dimer of pA-Tn5 fractions was pooled and concentrated by ultrafiltration to a final concentration of 60 μM and stored at −80°C until further use.

### in vitro reconstruction of Tn5 oligo complex

Preassembled Tn5 was prepared by following the protocols using Diagenode and a previous study^17^ with some modifications. Tagmentase (Tn5 transposase) - unloaded (Diagenode Cat# C01070010-10) was purchased from Diagenode and recombinant Tn5 protein was prepared in-house. Oligonucleotides (Tn5MErev, 5’-[phos] CTG TCTCTTATACACATCT-3’; Tn5ME-A, 5’-TCGTCGGCAGCGTCAGATGTGTATAAGAGACAG-3’; and Tn5ME-B, 5’-GTCTCGTGGGCTCGGAGATGTGTATAAGAGACAG-3’) were dissolved in TE buffer to prepare 100 μM stocks. Next, 1 μL of 10x annealing buffer (400 mM Tris-HCl, pH8.0, 500 mM NaCl) and 4.5 μL of 100 μM Tn5MErev was mixed with 4.5 μL of 100 μM Tn5ME-A or B and annealed using the following thermal cycler program: 95°C for 5 min, gradually cooling (−0.1°C/sec), 65°C for 5 min, gradually cooling to 4°C. The annealed oligos were mixed in one tube. An equal volume of annealed oligonucleotide mixture and Tn5 (18.7 μM) were mixed and incubated at 23°C for 30 min to assemble Tn5 and the annealed oligonucleotides. The preassembled Tn5 was diluted 10 times with 50% glycerol and stored at −20°C.

### Library preparation for sequencing

The Tn5 reaction solution was prepared by mixing 3.65 μL of water, 1.1 μL of 5x TAPS-DMF buffer (50 mM TAPS-NaOH, pH8.5, 25mM MgCl_2_, and 40% DMF), 0.25 μL of preassembled Tn5, and 0.5 μL of plasmid. The Tn5-based tagmentation reaction was performed by incubating at 55°C for 7 min. After adding 1.25 μL of 0.2% SDS, the tubes were incubated at 55°C for 7 min. The PCR reaction consisted of 5 μL of 2x KOD One master mix (TOYOBO), 4 μL of water, 0.5 μL of Tn5 reaction product, and 0.5 μL of illumina combinatorial dual index primer mixture (5′-AATGATACGGCGACCACCGAGATCTACACNNNNNNNNTCGTCGGCAGCGTC-3′ and 5′-CAAGCAGAAGACGGCATACGAGATNNNNNNNNGTCTCGTGGGCTCGG-3′, where NNNNNNNN means appropriate index). The PCR reaction was performed at 98°C for 2 min, followed by 30 cycles at 98°C for 10 s, 55°C for 5 s, and 68°C for 5 s. An equal volume of the PCR products was mixed in a single tube. Amplicons were separated by electrophoresis (100 V, 30–40 min) on a 3% agarose gel and 250–500 bp fragments were collected by gel purification. The concentration of the purified DNA libraries was measured using a Qubit3.0 fluorometer (Thermo Fisher Scientific) and diluted to 50 pM based on our previous report^18^. The 50 pM library was mixed with a 20–30% volume of 50 pM PhiX (Illumina) and sequenced using an iSeq100 (Illumina) in 150 bp-PE mode.

### Plasmid sequencing of E. coli lysates

A single colony was suspended in 10 μL water and lysed by incubating at 95°C for 10 min. After vortexing, the pellet was collected by centrifugation and 5 μL of the supernatant was removed from the remaining suspension. The *E. coli* genome was digested in the supernatant and the suspension with 0-10 U of exonuclease V (RecBCD) (New England Biolabs). The reaction was carried out in a 10-μL volume at 37°C for 1 h. EDTA (10 mM in final concentration) was added and ExoV was heat inactivated at 70°C for 30 min. The Tn5-based tagmentation reaction was done using 0.5 μL of the reaction product as indicated above.

### Bioinformatic pipeline for plasmid assembly

We implemented a script to encompass several key analysis tools and steps, each contributing to the overall process of automating the *de novo* assembly of plasmid, mapping to reference, and subsequent analysis. The script begins with a definition of the two essential functions that are responsible for handling plasmid assembly with or without providing reference sequence data. Using command line arguments, the script designates input and output directories, reference databases, and the input samplesheet file. Next, it scans the command line parameters to determine the input.zip file and the text file that contains relevant analysis details. To run the script, users must prepare their input data and execute the script in a Linux-based environment. This involves establishing a directory to organize the input data, placing FASTQ files and a text file with the analysis information in this directory, modifying the settings to match their system, and running the script via the terminal. The FASTQ files (read) in the directory are named according to the information in the text file used for the input. The information in the text file (.txt) contains information regarding user analysis. Each line of the text file represents a different analysis (sample) and should include the following columns (Tab-separated values):

1. Primary: The attributes used to identify a record.
2. read1: The name of the FASTQ file (. fastq.gz) containing the first set of short reads in each pair.
3. read2: The name of the FASTQ file (. fastq.gz) containing the second set of short reads in each pair.
4. expected_reads: The expected number of reads obtained from DNA sequencing.
5. plasmid_name: The name used for labeling the plasmid assembly data or designing the name of the plasmid.
6. output_directory: The name of the folder or directory where the results, data, or files generated from an analysis are stored.
7. Type_of_run: To specify the running of the analysis pipeline, using “y” to refer to running the plasmid assembly function by using a reference sequence for assembly, or using “n” to refer to the running the plasmid assembly pipeline with the *de novo* assembly workflow.
8. Reference genome filename: The name of the reference file in FASTA format (.fasta), which is required only if the ‘type_of_run’ is set to ‘y’; otherwise, it should be left empty.

The script iterates through each line of the text file, extracting the relevant information necessary for processing. Depending on the value specified in the “Type_of_run” column, the script follows one of two designed functions: The first function (Type_of_run = y) was designed for plasmid sequencing with a reference sequence. It uses a reference sequence for mapping. The mapping process includes the following steps: 1) Quality trimming of the reads by using fastp^19^; 2) the alignment of reads to the reference using Bowtie2^13^; 3) the resulting SAM file is converted to the sorted BAM format (.bam) using samtools^14^; 4) the coverage information is generated to describe the average number of reads that aligned to a known reference at specific locations within the target reference sequence; 5) Low-coverage regions were identified using BEDtools^15^ to filter these areas, as they may have higher error rates or may not be accurately represented in the assembled genome, being defined as ‘N’ in the consensus sequence; 6) a VCF file is created to capture mutations in the mapped sequence using bcftools^14^; and 7) The consensus sequence is generated from the VCF file using bcftools and the custom python script. Finally, a report file in “.html” format is generated using an in-house python script. The report includes the coverage information, and a mutation plot derived from the mutations in the VCF file. The consensus sequence serves as input for plasmid annotation, providing an annotation map in GenBank format and an interactive plasmid map in “.html” format via the pLannotate tool^16^.

The second function (Type_of_run = n) was generated for plasmid assembly through the *de novo* assembly of short sequencing reads. It was designed to streamline the data analysis without a reference sequence. The first step of the function involves applying fastp^18^ to assess the quality of the input files to eliminate low-quality reads as the first function. The *de novo* assembly of short sequencing reads was done using Unicycler tools^12^. A Burrows–Wheeler Alignment tool (BWA)^20^ was used for mapping by aligning the trimmed read sequences against the assembly sequence results from Unicycler. Finally, the assembled sequence serves as an input for plasmid annotation. It provides an annotation map in a GenBank format as well as an interactive plasmid map in .html format using the pLannotate tool as the method in the final mapping step^16^.

The script continues by generating individual assembly reports for each process run and indicates the performance of specific files, such as consensus sequences or annotations. A summary of the entire assembly process, encompassing a visualization of the various types of runs and their corresponding outcomes. This HTML report includes relevant statistics and graphical representations that enable users to ascertain the success of the assembly process.

## Reproducibility

A frozen version of the analysis script is available as a self-contained environment in a Docker hub (https://hub.docker.com/r/pgrrg/aistseq_analysis). The script used for the data analyses and source data is available on Github (https://github.com/aist-pgrrg/aistseq).

## Acknowledgments

We thank Y. Taguchi and S. Tsujikawa for their technical assistance. We also thank Dr. H. Tamaki for the use of the iSeq100 instrument. This work was supported by Japan Science and Technology Agency (JST) Moonshot R&D JPMJMS2217-2-3, and partly supported by JST-Core Research for Evolutional Science and Technology (CREST) JPMJCR19S6.

## Author contributions

S.S.S. conceived and designed the research. A.N. designed pA-Tn5 protein expression construct and performed most of the protein purification experiments. S.S.S., H.S., and K.M. prepared the library and performed sequencing. S.S.S., H.S., and J.R. conducted the data analysis. S.S.S. and J.R. built the pipeline for the sequencing analysis. H.S., K.M., J.R., and S.S.S. wrote the manuscript with the input of all the authors.

## Additional information

Supplementary information is available online.

