## Supplementary Figure S1 for "AistSeq: An In-House Tn5-Based Plasmid Sequencing Platform Using A Compact Benchtop Sequencer"

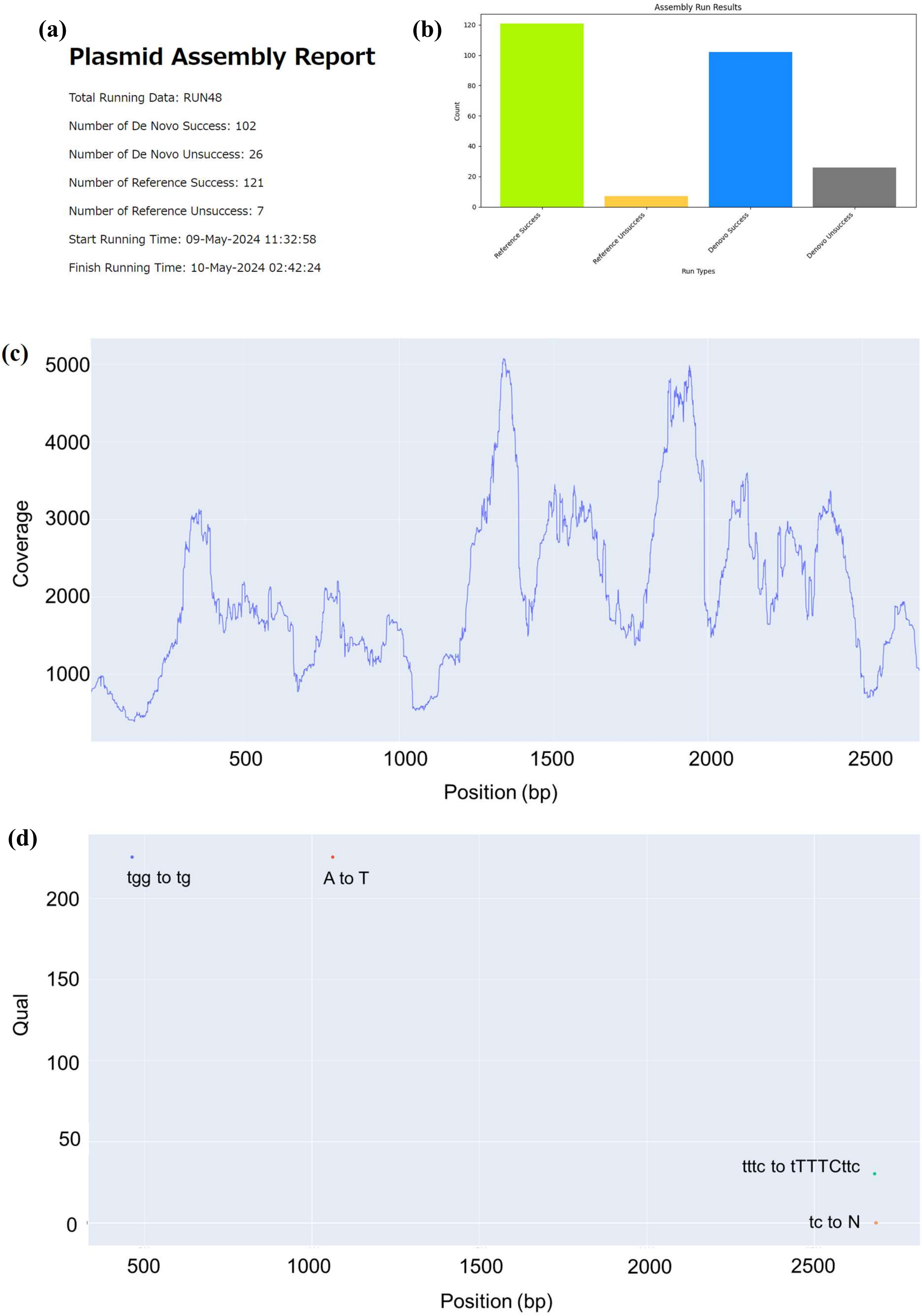
(a)
Plasmid Assembly Report
Total Running Data: RUN48
Number of De Novo Success: 102
Number of De Novo Unsuccess: 26
Number of Reference Success: 121
Number of Reference Unsuccess: 7
Start Running Time: 09-May-2024 11:32:58
Finish Running Time: 10-May-2024 02:42:24
(b)
Assembly Run Results
Count
0
20
40
60
80
100
120
Reference Success
Reference Unsuccess
Denovo Success
Denovo Unsuccess
Run Types
(c)
Coverage
1000
2000
3000
4000
5000
500
1000
1500
2000
2500
Position (bp)
(d)
Qual
0
50
100
150
200
tgg to tg
A to T
tttc to tTTTCttc
tc to N
500
1000
1500
2000
2500
Position (bp)

**(e)**

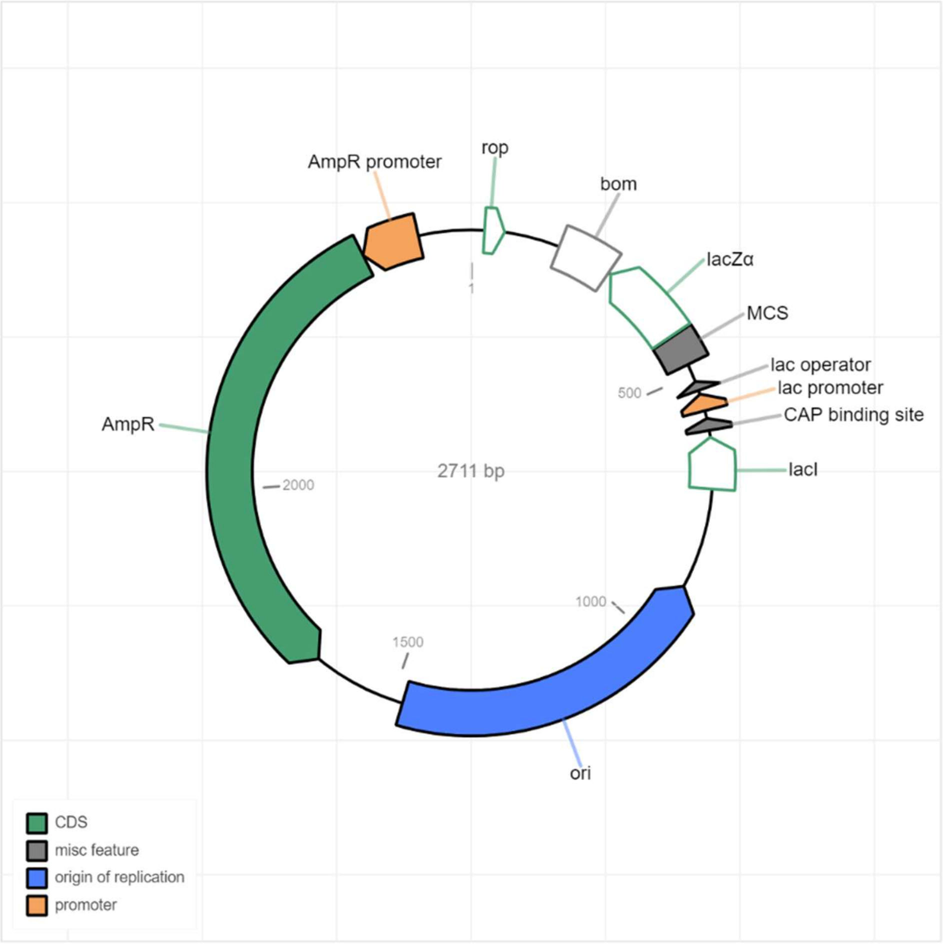


**Figure S1. Output data of the pipeline** (a) Representative image of the report of mapping to a reference and *de novo* assembly. (b) Representative coverage plot of a sample with low sequencing coverage based on the data from Table S1. (c) Representative coverage plot of a sample with sufficient sequencing coverage based on the data from Table S1. (d) Representative mutation plot for the pUC18 plasmid analyzed in this study. (e) The Representative output of an annotated plasmid map.
